# Genome sequencing for early-onset dementia: high diagnostic yield and frequent observation of multiple contributory alleles

**DOI:** 10.1101/748046

**Authors:** J. Nicholas Cochran, Emily C. McKinley, Meagan Cochran, Michelle D. Amaral, Bryan A. Moyers, Brittany N. Lasseigne, David E. Gray, James M.J. Lawlor, Jeremy W. Prokop, Ethan G. Geier, James M. Holt, Michelle L. Thompson, J. Scott Newberry, Jennifer S. Yokoyama, Elizabeth A. Worthey, David S. Geldmacher, Marissa Natelson Love, Gregory M. Cooper, Richard M. Myers, Erik D. Roberson

**Affiliations:** HudsonAlpha Institute for Biotechnology, 601 Genome Way, Huntsville AL 35806, USA; Alzheimer’s Disease Center, Department of Neurology, University of Alabama at Birmingham, 1720 7th Avenue South, Room 640, Birmingham AL 35294, USA; Department of Pediatrics and Human Development, Michigan State University, East Lansing, MI, USA; Department of Neurology, Memory and Aging Center, University of California San Francisco, 675 Nelson Rising Lane, Suite 190, San Francisco CA 94158, USA

**Author notes:** Richard M. Myers and Erik D. Roberson contributed equally to this study.

## Abstract

We assessed the utility of genome sequencing for early-onset dementia. Participants were selected from a memory disorders clinic. Genome sequencing was performed along with *C9orf72* repeat expansion testing. All returned sequencing results were Sanger validated clinically. Prior clinical diagnoses included Alzheimer’s disease, frontotemporal dementia, and unspecified dementia. The mean age-of-onset was 54 (41–76). 50% of patients had a strong family history, 37.5% had some, and 12.5% had no known family history. Nine of 32 patients (28%) had a variant defined as pathogenic or likely pathogenic (P/LP) by American College of Medical Genetics standards, including variants in *APP*, *C9orf72*, *CSF1R*, and *MAPT*. Nine patients (including three with P/LP variants) harbored established risk alleles with moderate penetrance (odds ratios of about 2–5) in *ABCA7*, *AKAP9*, *GBA*, *PLD3*, *SORL1*, and *TREM2*. All six patients harboring these moderate penetrance variants but not P/LP variants also had one or two *APOE* ε4 alleles. One patient had two *APOE* ε4 alleles with no other established contributors. In total, 16 patients (50%) harbored one or more genetic variants likely to explain symptoms. We identified variants of uncertain significance (VUSs) in *ABI3, ADAM10, ARSA, GRID2IP, MME, NOTCH3, PLCD1, PSEN1, TM2D3, TNK1, TTC3, and VPS13C*, also often along with other variants. In summary, genome sequencing for early-onset dementia demonstrated high utility, with particular advantages where targeted testing may fail such as atypical variant-disease associations or presence of multiple moderate impact alleles. One or more established contributory alleles is often present in early-onset dementia, supporting an oligogenic model.

## INTRODUCTION

Genomic technologies are increasingly being used in clinical settings, but clinical large-scale sequencing for adult-onset neurological conditions has not been heavily applied. Possible reasons include the use of disease-specific gene panels and uncertain genetic yield, despite promising signals for yield using comprehensive approaches (Blauwendraat et al. 2018). We sought to assess the diagnostic yield with genome sequencing and *C9orf72* expansion testing in cases of early-onset dementia.

Patients were selected from the Memory Disorders Clinic at the University of Alabama at Birmingham (UAB). Inclusion criteria were clinician-diagnosed early-onset dementia. When possible, unaffected parents were included as participants to allow filtering for *de novo* variants in patients without a family history (a fruitful approach in pediatric genetic disorders (Vissers et al. 2010; Bowling et al. 2017) and amyotrophic lateral sclerosis (ALS) (Chesi et al. 2013; Steinberg et al. 2015a)). In addition, unaffected siblings past the age of onset of the patient were enrolled as participants when possible for variant filtering and segregation.

Before starting analysis, we set criteria for return of results to patients. First, we used the American College of Medical Genetics (ACMG) criteria for pathogenicity (Richards et al. 2015) to identify highly penetrant causal variation. For moderately penetrant variants, we set criteria to return: (i) *APOE* ε4 status for early-onset Alzheimer’s disease (EOAD), (ii) any variant with a disease-associated odds ratio greater than two in multiple reports as an “established risk variant,” or (iii) one strong report with a disease-associated odds ratio greater than two with replication included in the study design as a “likely risk variant.”

## RESULTS

### Clinical presentation and family history

Prior clinical diagnoses for patients included EOAD, frontotemporal dementia (FTD), and other unspecified dementias. 21 patients were female and 11 were male. 28 self-reported Caucasian and four self-reported African American, all reported non-Hispanic ethnicity. The mean age of onset was 54 (range 41–76). 10 patients had ages of onset in their 40’s, 17 in their 50’s, 4 in their 60’s, and 1 in their 70’s.

In addition to enrolling patients, we also enrolled reportedly unaffected family members for variant filtering and segregation analyses. 31 unaffected relatives were enrolled, 29 of which had genome sequencing (2 were only checked for variants by Sanger). Only two families had complete trios (mother, father, and proband) to allow for searching for *de novo* variants, of which none of interest were identified. In total, 20 unaffected siblings, 9 unaffected parents, and 2 unaffected cousins were enrolled.

A strong family history of dementia was reported for 50% of patients (16/32), while 37.5% (12/32) had some family history, and 12.5% (4/32) had no reported family history. Our definition of family history is based on a modification of a four point scoring system first put forward by Jill Goldman (Goldman et al. 2005) where we modified the score as follows: (1) At least three people in two generations affected with EOAD, FTLD, ALS, CBD, or PSP with one person being a first-degree relative of the other two, (1.5) Same as (1) but with LOAD instead of EOAD, (2) At least three relatives with dementia or ALS but where criteria for autosomal dominant inheritance were not met, (3) A single affected first or second degree family member with early-onset dementia or ALS, (3.5) A single affected first or second degree family member with late-onset dementia or ALS, (4) No contributory family history or unknown family history. We considered a score of 1 or 1.5 as strong family history, a score of 2, 3, or 3.5 as some family history, and a score of 4 as no reported family history. All family history information is listed alongside phenotype and variant information in **Supplemental Table 1**.

To protect patient information, more detailed diagnoses and phenotype information beyond that provided here and listed in **Supplemental Table 1** are only provided in the controlled access dataset, NIAGADS project NG00082, to qualified researchers approved for access.

### Genomic analyses

Nine of 32 (28%) patients had a highly penetrant variant relevant to their clinical diagnosis (ACMG P/LP (Richards et al. 2015)), while seven (22%) had multiple moderately penetrant risk alleles (**Figure 1**). Individual cases are discussed next, with variants identified summarized by **Table 1** and listed alongside phenotype information in **Supplemental Table 1**.

**Figure 1.**
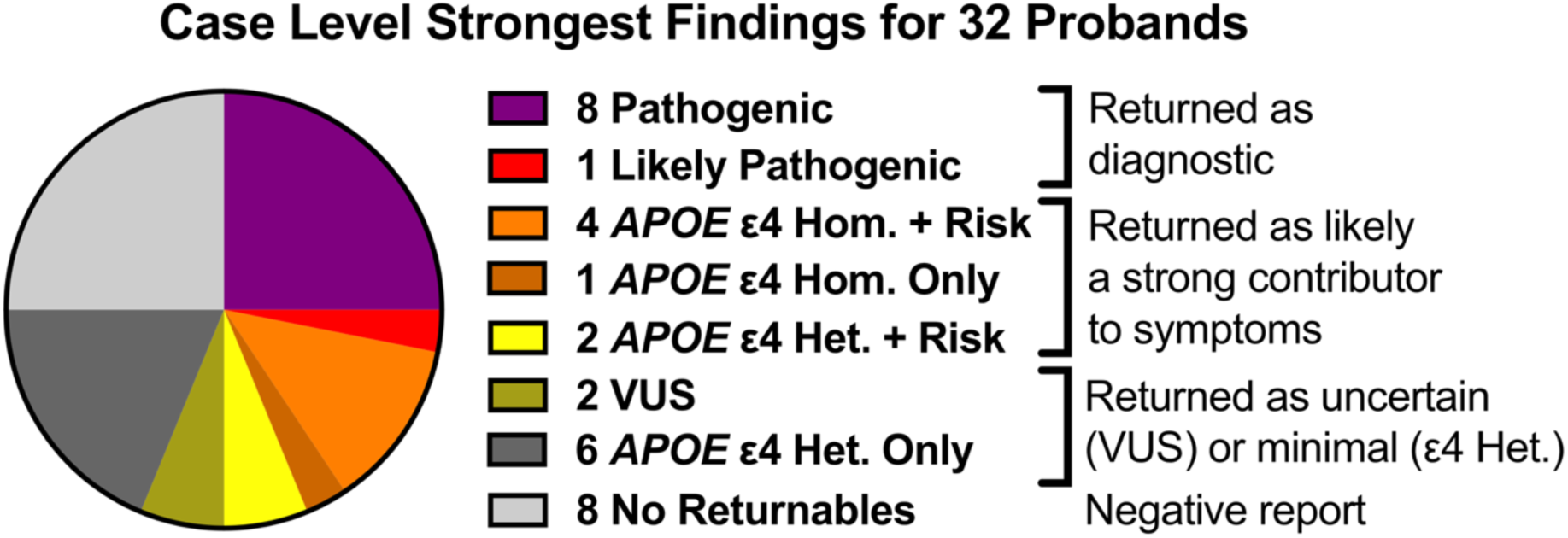
Summary of genomic analysis results for 32 patients with early-onset or familial dementia. Pathogenic variants were observed in APP (x2), C9orf72 (x3), and MAPT (x3). A likely pathogenic variant was observed in CSF1R. Four patients were APOE ε4 homozygous, with three of these patients also harboring additional risk variants in GBA, PLD3, and TREM2. Three patients were APOE ε4 heterozygous and had additional risk variants in AKAP9, SORL1, and TREM2. Two patients had variants of uncertain significance (VUS) in MAPT and NOTCH3. For six patients, the only returnable finding was APOE ε4 heterozygosity. Eight patients had no returnable findings.

**Table 1:**
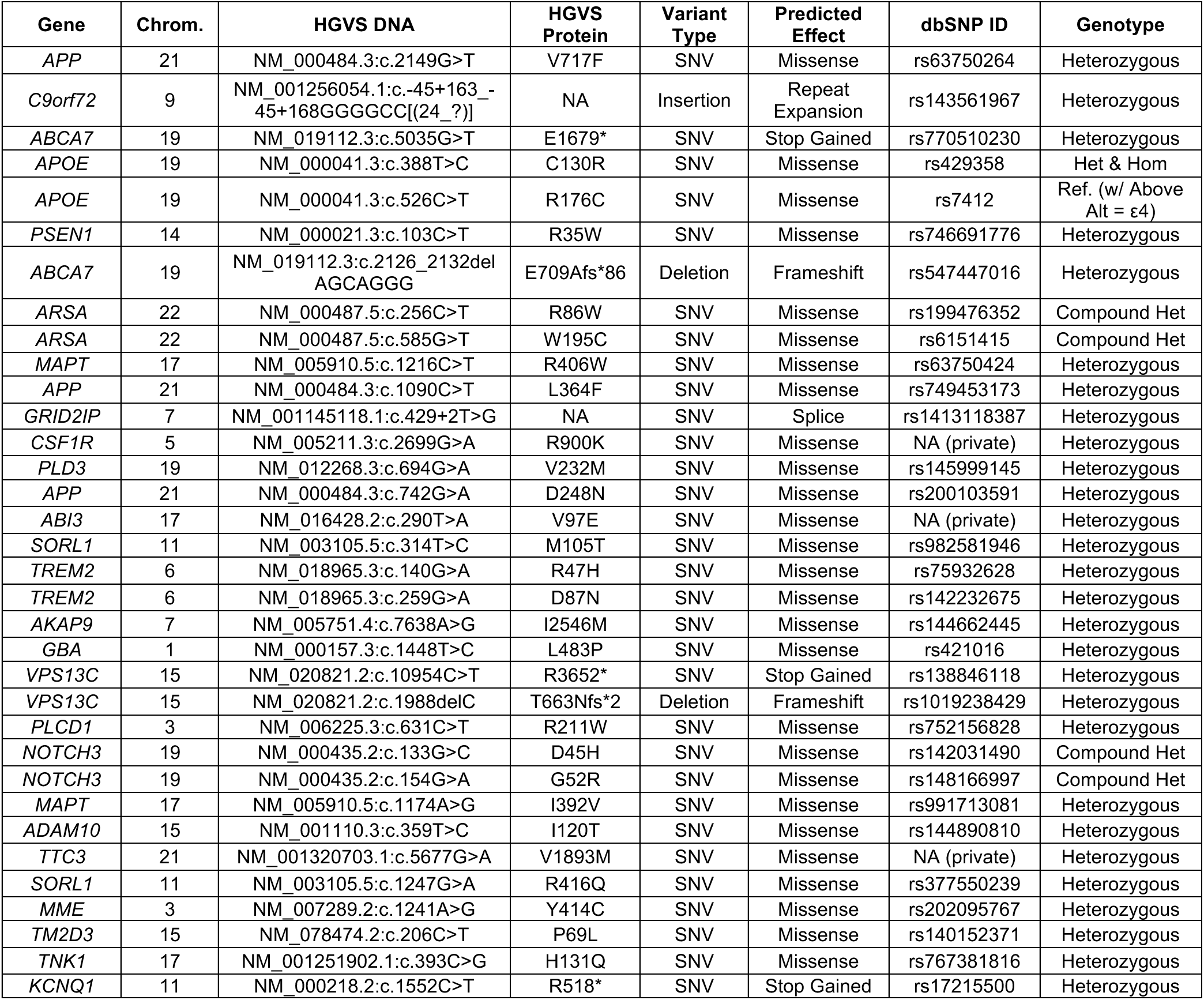
Variant Table. Note that many individuals had multiple candidate contributory variants, which is not captured when considering variants individually. For an expanded table that indicates multiple candidate variants, see **Supplemental Table 1**.

### Pathogenic or Likely Pathogenic Diagnoses

Variants were first evaluated using ACMG criteria for pathogenicity, and all P/LP variants were returned to patients (Richards et al. 2015). We provide a summary below, with detail on the ACMG evidence codes for variants provided in the Supplemental ACMG Pathogenicity Evidence Details.

### APP Pathogenic Variant (V717F) in Two Siblings

Two siblings with ages of onset in the mid-to-late 40s and a family history of EOAD suggestive of dominant inheritance harbored a pathogenic variant in *APP* (NM_000484.3, c.2149G>T, V717F), a well-established pathogenic variant (see Supplemental ACMG Pathogenicity Evidence Details). This variant is an example of one that would have been identified on commonly-used panels for genetic testing for EOAD.

### C9orf72 Expansion Carriers

Testing for a pathogenic G_4_C_2_ hexanucleotide expansion at the *C9orf72* locus associated with ALS and FTD was ordered for 30 of 32 patients (with two excluded for technical reasons, see Methods). GeneDx conducted a repeat-primed PCR test with 95% sensitivity and 98% specificity (Akimoto et al. 2014) to detect *C9orf72* expansions. As a technical aside, *C9orf72* expansions were not detectable using ExpansionHunter (Dolzhenko et al. 2017) or STRetch (Dashnow et al. 2018) in genome sequencing libraries prepared with PCR amplification assessed here. ExpansionHunter detects *C9orf72* expansions in PCR-free genome preparations (Dolzhenko et al. 2017), so PCR-free genome preparations or secondary testing (such as testing conducted by GeneDx for here) is necessary for detection of *C9orf72* expansions (and would also be necessary for other repeat expansions). Three patients with FTD (one patient also had ALS signs) with ages-of-onset in the 40s and 50s harbored a pathogenic expansion in *C9orf72* (see Supplemental ACMG Pathogenicity Evidence Details).

Some studies have suggested that additional contributing alleles could lower age of onset and/or alter clinical presentation for *C9orf72* expansion carriers (van Blitterswijk et al. 2012; van Blitterswijk et al. 2014; Pottier et al. 2015; Giannoccaro et al. 2017; Farhan et al. 2018). Consistent with this, all three *C9orf72* expansion carriers harbored other possibly contributory variants.

One carrier had three additional variants that may be contributory: an “established risk” stop gained variant in *ABCA7* (NM_019112.3, c.5035G>T, p.E1679*), one *APOE* ε4 allele, and a VUS in *PSEN1* (NM_000021.3, c.103C>T, p.R35W) (see Supplemental ACMG Pathogenicity Evidence Details). These variants may have contributed to the patient’s family history of multiple neurodegenerative diseases including ALS and EOAD.

Another carrier had a different “established risk” variant in *ABCA7* (NM_019112.3, c.2126_2132delAGCAGGG, p.E709Afs*86) (see Supplemental ACMG Pathogenicity Evidence Details), along with memory symptoms and a family history of AD, consistent with a possible contributory role of *ABCA7*.

The third carrier had two VUS in *ARSA*, associated with recessive metachromatic leukodystrophy (discussed further in Supplemental ACMG Pathogenicity Evidence Details).

### MAPT R406W Pathogenic Variant in Three Alzheimer’s Disease Patients

Three patients with EOAD (one patient also exhibited FTD signs) with ages-of-onset in the mid 50s to early 60s harbored a pathogenic variant in *MAPT* (NM_005910.5, c.1216C>T, p.R406W). Although *MAPT* pathogenic variants are typically associated with FTD (Cruts et al. 2012), this variant has been reported in patients with clinically diagnosed Alzheimer’s disease (AD) in multiple studies (see Supplemental ACMG Pathogenicity Evidence Details). This variant would not have been detected on many AD-specific panels, which often test for only *APP*, *PSEN1*, and *PSEN2*.

All three of these patients exhibited a possible contribution from another allele, just as in *C9orf72* expansion carriers. One patient had a loss-of-function “established risk” variant in *ABCA7* (NM_019112.3, c.2126_2132delAGCAGGG, p.E709Afs*86). Another patient had a VUS in *APP* (NM_000484.3, c.1090C>T, p.L364F). The third patient had a loss-of-function splice variant in *GRID2IP* (NM_001145118.1, c.429+2T>G), which, while not yet firmly associated with EOAD and thus not yet returnable, was implicated in a recent large sequencing study (Raghavan et al. 2018).

The presence of this rare variant in three individuals enrolled at the same clinic suggests they may share a common ancestor. However, none of these individuals are aware of any extended family members participating in the study. Furthermore, the patients are not detectably related by software used for routine checks of close familial relationships (KING).

### CSF1R R900K in an FTD Patient

A patient presenting with behavioral variant FTD (bvFTD) harbored a likely pathogenic variant in *CSF1R* (NM_005211.3, c.2699G>A, p.R900K) (see Supplemental ACMG Pathogenicity Evidence Details). Patients with variants in *CSF1R* can present with bvFTD, but the underlying pathology of pathogenic *CSF1R* variants is leukoencephalopathy (Rademakers et al. 2011; Stabile et al. 2016). Consistent with this, this patient had white matter abnormalities on MRI with frontal-predominant confluent white matter hyperintensity (**Figure 2A**) and global atrophy (**Figure 2B–D**). This variant would not have been detected on typical panels testing for FTD.

**Figure 2.**
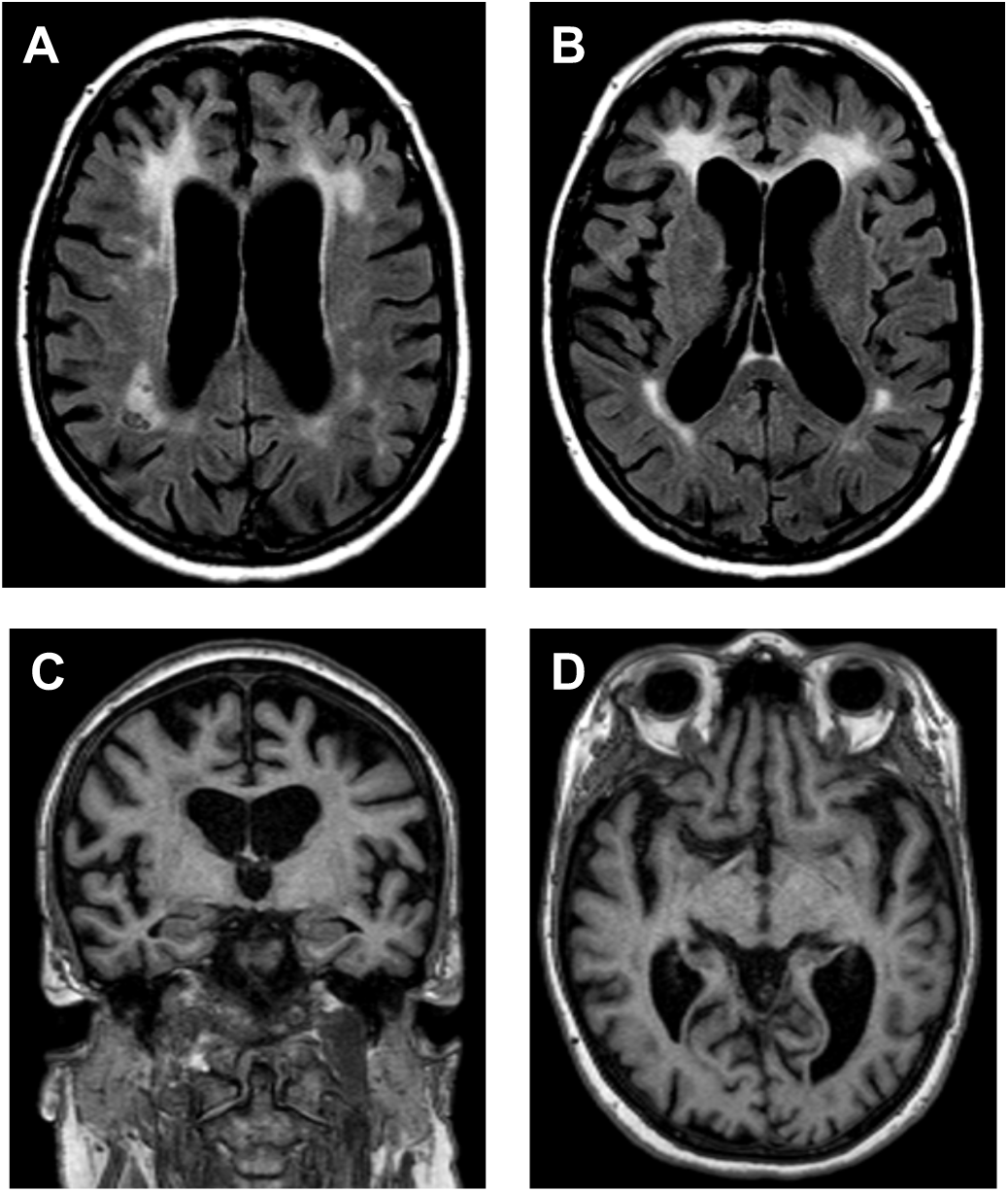
Neuroimaging findings in a CSF1R variant carrier. (**A,B**) Frontal-predominant, mildly asymmetric (R>L) white matter hyperintensities on axial FLAIR images. (**C,D**) Global cerebral atrophy on coronal and axial MPRAGE images. Radiological orientation with patient’s R side displayed on L.

### High Impact Risk Alleles

One unique aspect of this study is that we returned to patients moderately penetrant risk variants that meet criteria we have described. Intriguingly, rare variants meeting these criteria were observed only along with one or two *APOE* ε4 alleles, the most common moderately penetrant risk allele for AD (see Supplemental ACMG Pathogenicity Evidence Details). In all cases, *APOE* ε4 alleles were returned as “established risk variants.” The presence of one *APOE* ε4 allele was returned as likely only a small contributor to symptoms, while presence of two *APOE* ε4 alleles or one or two *APOE* ε4 alleles in combination with a rare moderately penetrant risk variant was returned with language indicating that such a combination of variants is likely to explain a large portion of the genetic contribution to symptoms (but with the caveat that family members should not be presymptomatically tested given incomplete penetrance). We continue with detail on some cases falling into this category.

### A case with APOE ε4 Homozygosity, PLD3 V232M, APP D248N, and ABI3 V97E

In a patient with EOAD whose symptoms began in the late 40s with enrolled unaffected parents, we observed an example of how EOAD may occur from a combination of inherited alleles from each parent, consistent with previous observations that EOAD can appear recessive in nature (Wingo et al. 2012). The patient had two *APOE* ε4 alleles (returned as “established risk,”) a *PLD3* variant (NM_012268.3, c.694G>A, p.V232M) (returned as “likely risk,”) an *APP* variant (NM_000484.3, c.742G>A, p.D248N) (returned as a VUS), and a private variant in *ABI3* (NM_016428.2, c.290T>A, p.V97E) (not returned but predicted damaging by PolyPhen-2 (Adzhubei et al. 2010) and SIFT (Ng and Henikoff 2003), with a CADD score (Kircher et al. 2014) of 33) (see Supplemental ACMG Pathogenicity Evidence Details). The *ABI3* variant was not returned to the patient because of insufficient evidence to consider the variant as a returnable VUS or risk variant, but is highlighted because a different coding variant in *ABI3* (NM_012268.3, c.1124T>C, p.S209F) (Sims et al. 2017) was associated with AD in a rigorous case-control study with an odds ratio of 1.4, yet is not predicted to be as damaging (CADD=13.5) and is relatively common in population databases (allele frequency of 0.6%). Therefore, we speculate that perhaps the variant we observed could have an effect of similar or greater magnitude given its higher predicted deleteriousness and absence from population databases. One of the *APOE* ε4 alleles and the variants in *PLD3* and *APP* was inherited from a parent with neurologic symptoms but not EOAD. The other parent harbored an *APOE* ε4 allele and the *ABI3* variant and did not have neurologic symptoms. This case serves as an example of how EOAD may arise with either no family history or limited family history of late-onset disease.

### A case with APOE ε4 Heterozygosity and SORL1 M105T

An individual with EOAD with onset in the mid 50s and a strong family history of AD had one *APOE* ε4 allele and a variant in *SORL1* (NM_003105.5, c.314T>C, p.M105T). While *SORL1* variants are not completely penetrant, loss-of-function variants in *SORL1* confer one of the highest levels of risk for AD outside of dominant pathogenic variants and *APOE*. Loss-of-function *SORL1* variant carriers in cases from a recent study (Raghavan et al. 2018) are present at an odds ratio of about four compared to population databases, a likely underestimate given that some individuals in population databases may develop AD. Indeed, a recent meta-analysis suggests the odds ratio for loss-of-function *SORL1* variants could be as high as 12.3 for all AD and 27.5 for EOAD (Campion et al. 2019).

For the *SORL1* variant identified here, we checked independent datasets for replication, and observed one M105T carrier in one study (Sassi et al. 2016), three M105T carriers in Alzheimer’s Disease Sequencing Project (ADSP) exomes (Bis et al. 2018), and two M105T carriers in ADSP genomes (one in an AD case and in one a mild cognitive impairment case) with no controls harboring the variant in any of these datasets. No other carriers were identified in cases or controls in four other studies (see Supplemental ACMG Pathogenicity Evidence Details). In addition to these four studies, there is one record in ClinVar from GeneDx (RCV000489328.1), but it lacked a denominator of the number of cases tested and thus was not considered in calculating the replication statistic. Taken together, *SORL1* M105T is observed six times out of 13,390 AD cases compared to 11 of 189,196 individuals at a population level for a replication-only odds ratio of 7.7 (*p* = 0.0005 by Fisher’s exact test). This variant did not completely segregate with disease in four family members of our patient. However, the age-of-onset range for similar variants in *SORL1* can be up to 24 years (Louwersheimer et al. 2017), which is wider than the age differences between the family members we genotyped, suggesting that this segregation analysis may not be completely informative. Considering all of the evidence, we returned this variant to the patient as a VUS (it could also be considered a “likely risk variant”). Modelling suggests M105T is a highly conserved residue (**Figure 3A**) where change to a threonine may create a PLK1 kinase site that may disrupt function (**Figure 3B**) (discussed further in Supplemental ACMG Pathogenicity Evidence Details).

**Figure 3.**
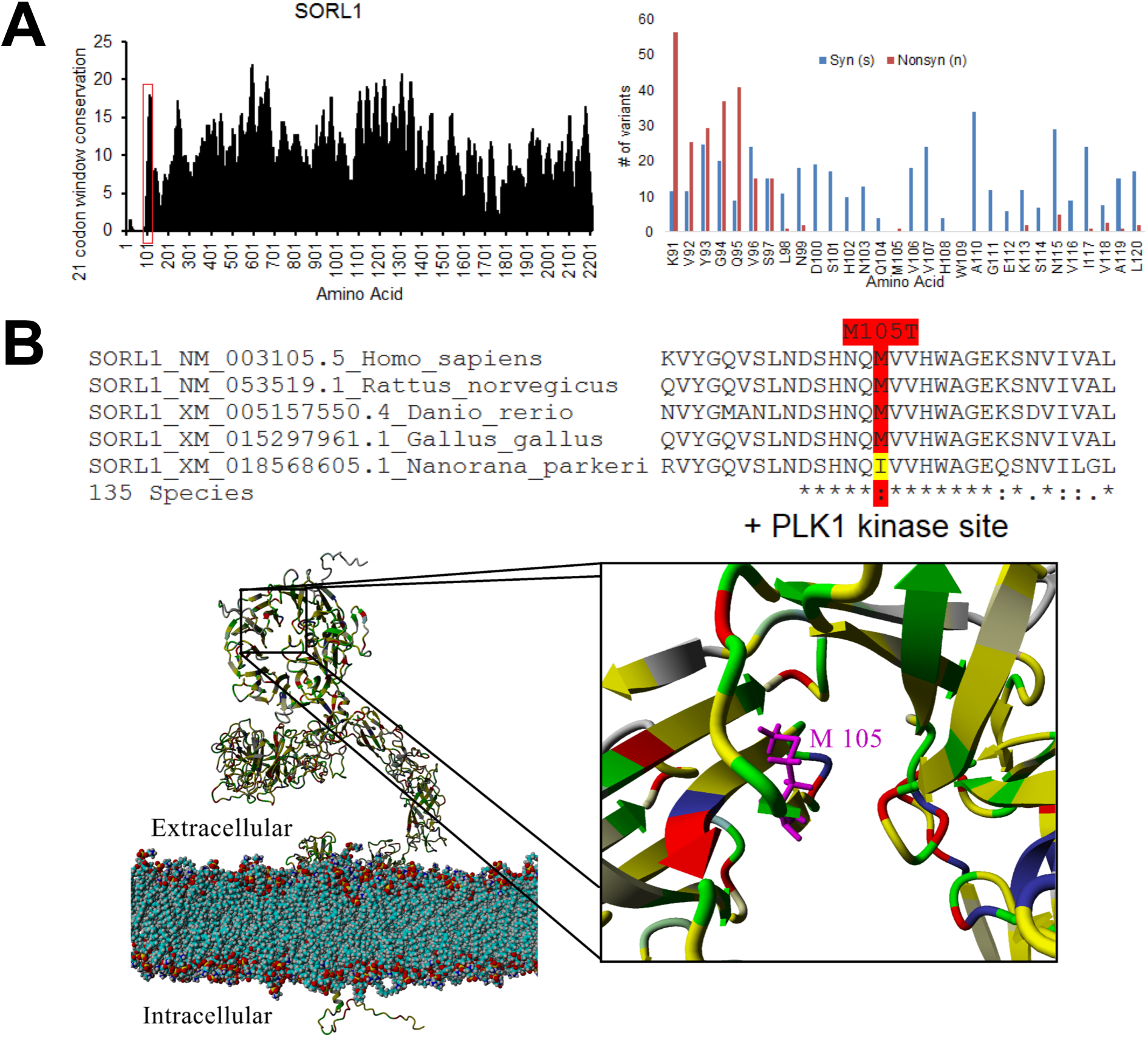
Molecular modeling of the effect of the M105T variant on SORL1. (**A**) Conservation analysis of the SORL1 gene sequence was performed across open reading frame sequences of 135 species. Scores at each codon were assessed with 100% conservation receiving a score of 1, with addition of a score for codon selection (score of 0 if dN-dS of site is below mean, 0.25 for sites with values above the mean to one standard deviation above the mean, 0.5 for sites greater than one standard deviation but below two standard deviations, one for sites greater than two standard deviations). A score of two is maximal, suggesting an amino acid that is 100% conserved with codon wobble indicative of a high selection rate at the position. The values were then placed on a 21-codon sliding window (combining values for 10 codons before and after each position) to identify conserved motifs within the gene. (**B**) Model of SORL1 protein (assessed with YASARA2). Colors are based on 135 species alignments fed into ConSurf such that colors indicate: gray=not conserved, yellow=conserved hydrophobic, red=conserved polar acidic, blue=conserved polar basic, green=conserved hydrophilic. Note that the M105T variant leads to a predicted gain of a PLK1 kinase target site in SORL1.

### APOE ε4 with TREM2, AKAP9, and GBA Risk Variants

In two cases with EOAD beginning in the late 40s, we observed a risk allele in *TREM2* and one or two *APOE* ε4 alleles. The first was *TREM2* (NM_018965.3, c.140G>A, p.R47H) (Guerreiro et al. 2013; Jonsson et al. 2013) with one *APOE* ε4 allele. This *TREM2* variant was returned as an “established risk variant.” Second, we observed *TREM2* (NM_018965.3, c.259G>A, p.D87N) (Guerreiro et al. 2013) (see Supplemental ACMG Pathogenicity Evidence Details) with two *APOE* ε4 alleles. This *TREM2* variant was returned as a “likely risk variant.”

In an African American patient with features of both EOAD and FTD, we observed a variant in *AKAP9* previously reported to increase risk in African Americans (NM_005751.4, c.7638A>G, p.I2546M) (Logue et al. 2014). In this case, despite only being observed in one study with replication, the specificity of this variant disease association to African American ethnicity and additional functional data (Ikezu et al. 2018) provided enough evidence to return this as an “established risk variant.”

A patient with EOAD with onset in the mid 50s harbored *GBA* (NM_000157.3, c.1448T>C, p.L483P [previous nomenclature, p.L444P]) and two *APOE ε4* alleles, originally associated with Lewy body disorders (Mata et al. 2008), but later also with mixed Dementia with Lewy Bodies and AD (Tsuang et al. 2012; Nalls et al. 2013). Because of this and a recent association with accelerated cognitive decline (Liu et al. 2016), we returned this as a “likely risk variant.”

### VPS13C loss-of-function with APOE ε4

A patient with mixed symptoms of AD and FTD with onset in the late 60s harbored *VPS13C* (NM_020821.2, c.10954C>T, p.R3652*) and two *APOE* ε4 alleles. A patient with EOAD with onset in the late 40s had *VPS13C* (NM_020821.2, c.1988delC, p.T663Nfs*2), a variant in *PLCD1* (NM_006225.3, c.631C>T, p.R211W), and one *APOE* ε4 allele. Only *APOE* ε4 was reported back to these patients because of uncertain contribution of the other variants to the phenotype. Homozygous loss of *VPS13C* is associated with early-onset Parkinson’s (Schormair et al. 2018). We do not know the significance of the observation of one loss-of-function allele here, although unpublished studies have reported an association between heterozygous loss-of-function in *VPS13C* and FTD (see Supplemental ACMG Pathogenicity Evidence Details). *PLCD1* was proposed as a candidate gene for AD in one study (Shimohama et al. 1998). Observing two loss-of-function variants in *VPS13C* in this small cohort leads us to speculate that heterozygous loss-of-function variants in *VPS13C* may contribute to early-onset dementia.

### Variants of Uncertain Significance or Research Interest

Five other patients harbored interesting – but speculative – VUSs or combinations of variants of interest for future research. These include (1) a patient with possible CADASIL and a haplotype of uncertain significance with two variants in *NOTCH3 (*NM_000435.2, c.133G>C, p.D45H and NM_000435.2, c.154G>A, p.G52R), (2) a patient with a VUS in *MAPT* (NM_005910.5, c.1174A>G, p.I392V), (3) a patient with an *APOE* ε4 allele and a variant in both *ADAM10* (NM_001110.3, c.359T>C, p.I120T) and *TTC3* (NM_001001894.2, c.5557G>A, p.V1853M), (4) a patient with an *APOE* ε4 allele, and a variant in both *SORL1* (NM_003105.5, c.1247G>A, p.R416Q) and *MME* (NM_007289.2, c.1241A>G, p.Y414C), and (5) a patient with variants in *TM2D3* (NM_078474.2, c.206C>T, p.P69L) and *TNK1* (NM_001251902.1, c.393C>G, p.H131Q). Furthermore, one patient harbored a secondary pathogenic variant in *KCNQ1* (NM_000218.2, c.1552C>T, R518*). We expand on these cases in the Supplemental ACMG Pathogenicity Evidence Details.

### Quantitative Enrichment of Multiple Alleles

Because we observed so many cases harboring multiple established alleles, we asked if this effect was statistically enriched over a control population recruited from the same geographical area, with controls reporting a family history of dementia excluded. We set criteria for qualifying variants as follows: (1) *TREM2* or *GBA* missense or loss-of-function variants with CADD>20 and population frequency <0.5% in both gnomAD (Lek et al. 2016) and TOPMed Bravo (NHLBI 2018), (2) *ABCA7*, *SORL1*, *TBK1*, or *GRN* loss-of-function variants with CADD>20 and population frequency <0.5%, (3) the specific *PLD3* and *AKAP9* variants observed here (since their associations are for single alleles), missense only variants with CADD>20 and population frequency <0.01% for *SORL1*, *CSF1R*, *APP*, *PSEN1*, *PSEN2*, and *MAPT*, (5) expansion carriers in *C9orf72*, and (6) *APOE* ε4 alleles. We recognize that this may contain bias since these filtering criteria were selected after analysis of cases. However, we attempted to mitigate this by selecting reasonable thresholds that would catch variants not identified in this study but that would still have been considered if they had been identified. For example, we did not observe any variants meeting these criteria in *TBK1* or *GRN* but included them here because of their important role in disease. We also included *C9orf72* carriers without information on if any are present in the control population, but this is a reasonable assumption (see Supplemental ACMG Pathogenicity Evidence Details).

Variants meeting the criteria described are highly enriched in cases (**Figure 4A**). Intriguingly, there is no enrichment of *APOE* ε4 alleles in the absence of other qualifying alleles (**Figure 4B**). In contrast, the presence of *APOE* ε4 alleles in combination with another qualifying variant is highly enriched in cases, regardless of whether Mendelian variants are included in the calculation (**Figure 4C**) or excluded (**Figure 4D**). The odds ratios for *APOE* ε4 alleles in combination with another qualifying variant in cases without a Mendelian cause suggests that the presence of rare variants increases odds ratios approximately multiplicatively over those typically reported for *APOE* ε4 alone (typically reported: ∼2.5 for one *APOE* ε4 allele, with a rare variant, 5.5; 10–15 for two *APOE* ε4 alleles, with a rare variant, 39.1), see Supplemental ACMG Pathogenicity Evidence Details on *APOE*) (**Figure 4D**).

**Figure 4.**
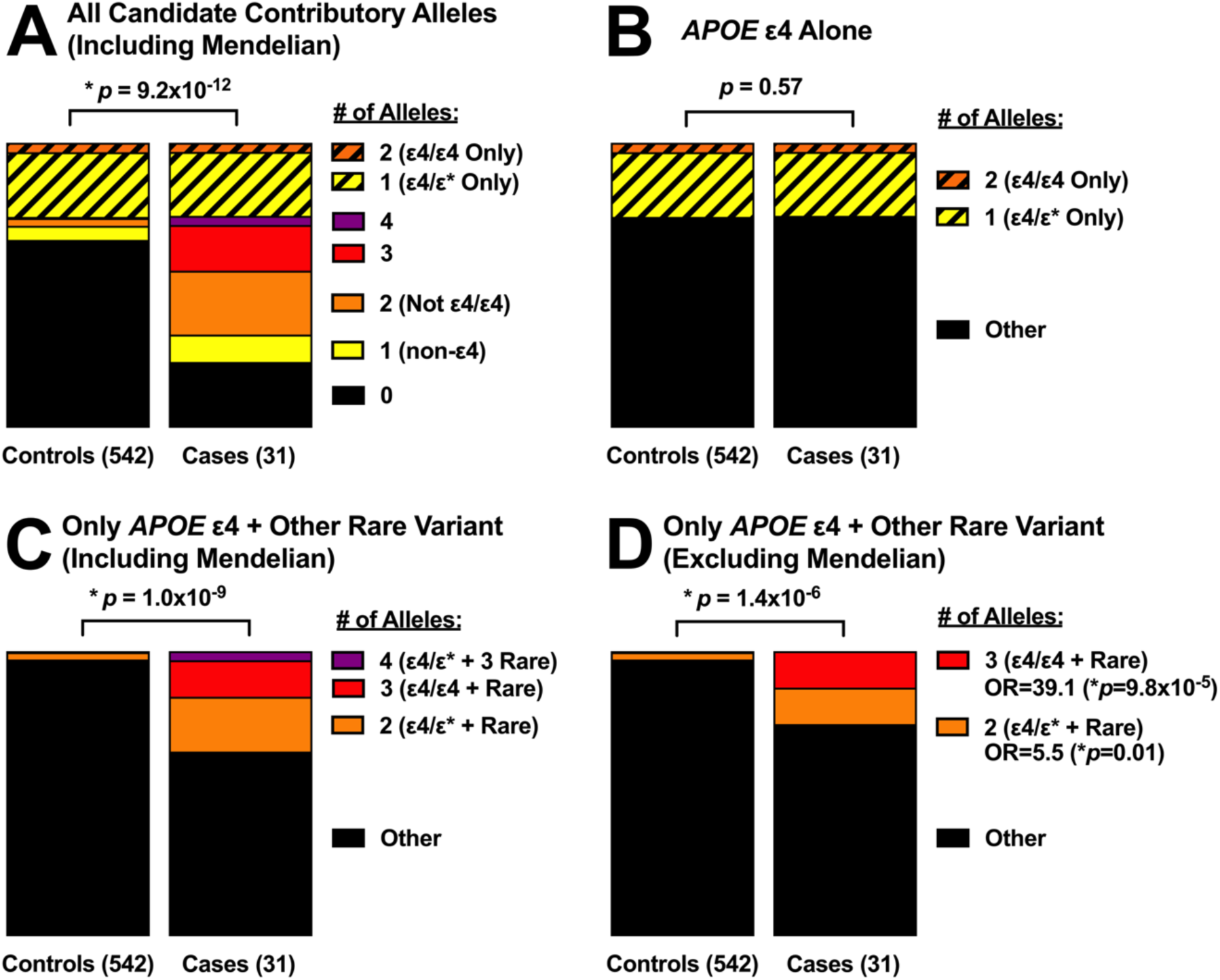
Multiple variants in neurodegeneration-associated genes are often observed in early-onset dementia, with a critical role for rare variants acting in combination with APOE ε4. Note: for all panels, ε4/ε* indicates either ε4/ε3 or ε4/ε2 (mostly ε4/ε3). Also for all panels, cases N=31 (32 probands excluding 1 sibling from an affected sibling pair) and controls N=542. (**A**) Qualifying candidate alleles associated with neurodegeneration (see text for criteria) are highly enriched in cases (p=9.2×10^-12^ by exact conditional Cochran-Armitage trend test). (**B**) Presence of APOE ε4 alone, in the absence of any other qualifying variants, is not enriched in cases (p=0.57 by exact conditional Cochran-Armitage trend test). (**C**) Presence of APOE ε4 along with at least one qualifying rare variant (including Mendelian variants) is highly enriched in cases (p=1.0×10^-9^ by exact conditional Cochran-Armitage trend test). (**D**) Presence of APOE ε4 along with at least one qualifying rare variant (excluding Mendelian variants) is highly enriched in cases (p=1.4×10^-6^ by exact conditional Cochran-Armitage trend test). The odds ratio for Presence of one APOE ε4 allele along with one qualifying rare variant vs. controls is 5.5 (p=0.01 by Fisher’s exact test, 95% CI 1.2–19.3). The odds ratio for Presence of two APOE ε4 alleles along with one qualifying rare variant vs. controls is 39.1 (p=9.8×10^-5^ by Fisher’s exact test, 95% CI 5.3–447.5).

## DISCUSSION

One key theme in this study was the frequent observation of multiple possible contributory alleles. We even observed this in multiple cases with clear, highly penetrant, pathogenic variants despite a small cohort size. The degree to which additional alleles contribute in dominant cases cannot be assessed without larger cohorts to evaluate effects on age-of-onset or other variables. However, given that other studies have made similar observations in ALS/FTD (van Blitterswijk et al. 2012; van Blitterswijk et al. 2014; Pottier et al. 2015; Giannoccaro et al. 2017; Farhan et al. 2018), this phenomenon clearly warrants further investigation.

In cases where a dominant pathogenic variant was not present, there was enrichment for multiple established alleles contributing to disease risk. Every case with a moderately penetrant risk variant established by case-control studies identified in this cohort also harbored one or two *APOE* ε4 alleles, emphasizing the importance of *APOE* ε4. Future efforts in analysis of large cohorts should include analysis of level of risk when rare risk variants are present, for example by incorporation of signal from rare variation in established risk genes into polygenic risk scores. Several groups have begun developing polygenic risk scores for AD (Escott-Price et al. 2015; Desikan et al. 2017), but these scores are based solely on common variation. This is, of course, a reasonable approach because it maximizes reproducibility, as considering rare variants could lead to an over-trained model. However, while rare variants are rare individually, aggregation approaches may provide replicable and meaningful signals if incorporated for key genes where rare variants are now established to confer risk for AD, such as *ABCA7*, *SORL1*, and *TREM2*. Similarly, while large FTD genetic studies are not as progressed as those for AD, we can begin to consider genes where variation in a polygenic risk score may be informative for FTD, such as *TBK1* (Cirulli et al. 2015), *MFSD8* (Geier et al. 2019), *DPP6*, *UNC13A*, and *HLA-DQA2* (Pottier et al. 2019).

In Conclusion, this study demonstrates the high diagnostic and research utility of genome sequencing in cases of early-onset dementia. Mendelian diagnostic yield in this population was 28%, with an additional 22% of patients harboring risk-increasing variants that, in combination with *APOE* ε4, likely account for most of the genetic contribution to their symptoms. Genome sequencing is able to identify relevant variation in conditions with high genetic heterogeneity, nonspecific phenotypes, or established risk factors that do not follow a clear Mendelian pattern, and allowed for identification of cryptic genotype-phenotype relationships that likely would have been missed by panel testing. In addition to the research value of this study, it had value for patient care as well, for example by allowing for referral of families to the Dominantly Inherited Alzheimer’s Network and the Advancing Research & Treatment for Frontotemporal Lobar Degeneration study. We conclude that application of more comprehensive genetic testing (including genome sequencing, where appropriate) could aid in evaluation of early-onset dementia cases currently and will continue to grow in utility for future use.

## METHODS

### Genome sequencing

Genome sequencing was performed at the HudsonAlpha Institute for Biotechnology on Illumina HiSeq X or NovaSeq platforms using paired end 150 base pair reads. Mean depth was 34X with an average of 91.5% of bases covered at 20X. Sequencing libraries were prepared by Covaris shearing, end repair, adapter ligation, and PCR using standard protocols. Library concentrations were normalized using KAPA qPCR prior to sequencing. All sequencing variants returned to patients were validated by CAP/CLIA Sanger.

### Data processing and quality control

Demuxed FASTQs were aligned with bwa-0.7.12 (Li and Durbin 2009) to hg19. BAMs were sorted and duplicates were marked with Sambamba 0.5.4 (Tarasov et al. 2015). Indels were realigned, bases were recalibrated, and gVCFs were generated with GATK 3.3 (McKenna et al. 2010). gVCFs were batch called with GATK 3.8. KING 2.1.2 (Manichaikul et al. 2010) was used for sex checks on VCFs, for validation of known familial relationships, and to check for unknown familial relationships (none of which were identified).

### C9orf72 expansion testing

Samples from 30 of 32 patients were tested for pathogenic *C9orf72* repeat expansion alleles by GeneDx (Gaithersburg, MD). Two patients did not have sufficient material for testing, but both lacked symptoms consistent with a *C9orf72* repeat expansion and also had another likely explanation of symptoms: one had a pathogenic *APP* variant and another harbored both one *APOE* ε4 allele and a *TREM2* established risk allele).

### Genomic data analysis

The HudsonAlpha-developed Codicem application (http://envisiongenomics.com/codicem-analysis-platform/) was used to analyze and support interpretation of the variant data (described elsewhere (Holt et al. 2019)). Although this software package was used for analysis, it would not be necessary to use this package to reproduce this work. Simple filtering for population allele frequencies (ie gnomAD (Lek et al. 2016) and TOPMed Bravo (NHLBI 2018)), *in silico* deleteriousness scores (ie CADD (Kircher et al. 2014), PolyPhen-2 (Adzhubei et al. 2010), and SIFT (Ng and Henikoff 2003)), and gene lists relevant to the phenotype of interest would recapitulate our findings using any suitable software package, or even by a command line interface.

In addition to searching for single nucleotide variants and small indels, we also searched for large copy number variations using four callers (DELLY (Rausch et al. 2012), ERDS (Zhu et al. 2012), CNVnator (Abyzov et al. 2011), and BIC-seq2 (Xi et al. 2016)), but did not identify any relevant to patient phenotypes (including absence of APP duplications).

### SORL1 structural modeling

SORL1 structural modeling and evolutionary conservation analysis was assessed using a previously published sequence-to-structure-to-function workflow (Prokop et al. 2017).

### Statistics

The exact conditional Cochran-Armitage trend test was calculated using the CATTexact 0.1.0 package and Fisher’s exact test using fisher.test in R 3.4.1.

### Return of results

Results meeting criteria for return were delivered to patients by clinicians in the UAB Memory Disorders Clinic through letters written by a genetic counselor. Letters included information on the variant, associated disease, recurrence risk, and management recommendations. Patients were given the option to have a genetic counselor present for return of results via phone or videoconference or to follow up with a genetic counselor after delivery of results. Primary results were provided only to probands. Although a secondary result was identified in only one participant who was a patient, we did also offer non-patient participants (family members) receipt of actionable secondary findings (ACMG 59™) if such a result had been identified. Family members of patients that received diagnostic results were provided with information to seek out clinical genetic counseling and targeted testing for familial variants if they desired.

## ADDITIONAL INFORMATION

### Data Deposition and Access

All data from participants enrolled as a part of this study, including more detailed phenotype data for the cases described here, are available on the National Institute on Aging Genetics of Alzheimer’s Disease Data Storage (NIAGADS) site under project NG00082. Data from control subjects not enrolled as a part of this study are available under dbGaP accession phs001089.v3.p1, which contains data generated by the Clinical Sequencing Exploratory Research (CSER) Consortium established by the NHGRI. Funding support for CSER was provided through cooperative agreements with the NHGRI and NCI through grant numbers U01 HG007301 (Genomic Diagnosis in Children with Developmental Delay). Information about CSER and the investigators and institutions who comprise the CSER consortium can be found at https://cser-consortium.org.

ADNI (Alzheimer’s Disease Neuroimaging Initiative, part of the ADSP genomes batch call) and ADSP data are available at NIAGADS under projects NG00066 and NG00067 and on dbGap under accession phs000572.v7.p4 (see Supplemental Extended Acknowledgements for full list of ADNI and ADSP contributors and funding sources).

### Ethics Statement

This study was approved by UAB IRB protocol X161202004, “Evaluation of Genomic Variants in Patients with Neurologic Diseases.” All participants described provided explicit written consent for publication.

## Acknowledgements

We thank Alissa Mackiewicz from the HudsonAlpha Foundation for assistance in securing funding, Jennifer Mahaffey at UAB for assistance with the IRB application, Mackenzie Fowler at UAB for assistance with participant recruitment, the Clinical Services Lab and the Genomic Services Lab at HudsonAlpha for DNA isolations, library generation, quality control and sequencing, the Codicem software development team at HudsonAlpha for genome analysis software, David Bick at HudsonAlpha for helpful discussions about ACMG guidelines, and Dominique Campion at University of Rouen for correspondence indicating the absence from both cases and controls of the M105T variant in *SORL1* in the dataset published in (Bellenguez et al. 2017).

## Authors’ contributions

JNC, GMC, RMM, and EDR designed the study. JNC and RMM secured funding. JNC and EDR wrote the IRB protocol. ECM coordinated all aspects of patient interaction. JNC, MDA, BAM, and BNL analyzed genomes with input from MEC, ECM, and EDR. MDA coordinated *C9orf72* testing. JNC, DEG, JMJL, JWP, EGG, JMH, and JSN conducted other analyses. MEC wrote clinical letters and provided genetic counseling. MLT provided phenotype information for controls. JSY accessed ADSP and supervised EGG. EAW supervised JMH, JSN, and the software development team. EDR, DSG and MNL recruited participants and returned results. GMC supervised DEG, JMJL, and MLT. JNC wrote the manuscript, with edits by ECM, MEC, MDA, BAM, BNL, JWP, EGG, JMH, EAW, GMC, and EDR. All authors approved the final manuscript.

## SUPPLEMENTAL MATERIAL

### ACMG Pathogenicity Evidence Details

#### *APP* (NM_000484.3, c.2149G>T, V717F)

Two strong criteria, three moderate criteria, and one supporting criterion result in the ACMG-recommended assertion of “pathogenic.”

- Strong segregation data (Murrell et al. 1991; Finckh et al. 2005) (ACMG criterion PP1S)
- Biochemical studies (Tamaoka et al. 1994; Nilsberth et al. 2001; Sato et al. 2003) (ACMG criterion PS3)
- The same amino acid is mutated to other amino acids by other segregating EOAD pathogenic variants (Chartier-Harlin et al. 1991; Goate et al. 1991; Murrell et al. 2000), and others reviewed in the AD&FTD Mutation Database (Cruts et al. 2012) (ACMG criterion PM5)
- The variant is located in a well-established functional domain at the epsilon cleavage site for gamma secretase (Dimitrov et al. 2013) (and reviewed in (Holtzman et al. 2011)) (ACMG criterion PM1)
- Absent from the gnomAD (Lek et al. 2016) and TOPMed Bravo population databases (NHLBI 2018) (ACMG criterion PM2)
- Predicted damaging by multiple computational methods (CADD (Kircher et al. 2014), PolyPhen-2 (Adzhubei et al. 2010), and SIFT (Ng and Henikoff 2003)) (ACMG criterion PP3).

#### C9orf72 Expansion Carriers

- Strong segregation with ALS and FTD (DeJesus-Hernandez et al. 2011; Renton et al. 2011) (ACMG criterion PP1S).
- Extensive functional studies support the pathogenicity of this allele (key examples in (Chew et al. 2015; Zhang et al. 2015), and recently reviewed in (Babic Leko et al. 2019; Vatsavayai et al. 2019)) (ACMG criterion PS3).
- *Note on the assumption that C9orf72 expansions will be absent from controls:* two studies have assessed the frequency of *C9orf72* expansions in healthy controls, both arriving at a frequency of approximately 0.2% of individuals (Beck et al. 2013; Kaivola et al. 2019) (this would be equivalent to approximately 1 carrier in our control set of 542 individuals). However, in one of these studies, they also assessed for other neurologic diseases, and found that 4 of 6 individuals with *C9orf72* expansions (out of 3142) had another neurologic disease (Kaivola et al. 2019), leaving only 2 expansion carriers out of 3142 individuals in that study. Therefore, the assumption that no repeat expansion carriers are present in the control set we selected where individuals with a family history of any neurologic disease have been excluded is not unreasonable.

#### ARSA alleles

In one *C9orf72* expansion carrier, we identified a possibly contributory combination of variants in *ARSA*, associated with recessive metachromatic leukodystrophy (which can include dementia as a symptom): one reported pathogenic variant that may maintain some residual activity (an “R” allele) (NM_000487.5, c.256C>T, p.R86W), and one variant of uncertain significance (VUS) (NM_000487.5, c.585G>T, p.W195C) that may be a pseudo-deficiency allele. Because we did not have phasing data for these two variants and could not follow up with a biochemical test of enzyme activity (the patient died between study enrollment and the observation of the variants in *ARSA*), the specific contribution of these variants is unknown.

- Reported pathogenic variant that may maintain some residual activity (an “R” allele) (NM_000487.5, c.256C>T, p.R86W) (Biffi et al. 2008; Cesani et al. 2016)
- Reported Variant of uncertain significance (VUS) (NM_000487.5, c.585G>T, p.W195C) that may be a pseudo-deficiency allele (Xiong et al. 2015; Cesani et al. 2016; Dehghan Manshadi et al. 2017)
- These alleles were reported together as a VUS, with special emphasis that this combination of alleles may have no or little influence on disease given the presence of a *C9orf72* expansion
- https://rarediseases.org/rare-diseases/metachromatic-leukodystrophy/

#### *ABCA7* Loss-of-Function Alleles

- We identified two loss-of-function variants in *ABCA7*: (NM_019112.3, c.2126_2132delAGCAGGG, p.E709Afs*86) and *ABCA7* (NM_019112.3, c.5035G>T, p.E1679*). Loss-of-function variants in *ABCA7* have been solidly associated with AD by several independent case-control studies (Cuyvers et al. 2015; Del-Aguila et al. 2015; Steinberg et al. 2015b; Allen et al. 2017; De Roeck et al. 2017; N’Songo et al. 2017).

#### *APOE* ε4 allele

- The APOE ε4 allele is definitively established by a plethora of studies to be associated with AD, with a few key references noted here (Corder et al. 1993; Saunders et al. 1993; Farrer et al. 1997; Lambert et al. 2013; Yu et al. 2014; Qian et al. 2017).

#### *PSEN1* (NM_000021.3, c.103C>T, p.R35W)

- Another VUS in *PSEN1* has been described at Arg35 that does not completely segregate with disease (Rogaeva et al. 2001; Raux et al. 2005; Benitez et al. 2013).

#### *MAPT* (NM_005910.5, c.1216C>T, p.R406W)

- Strong segregation with EOAD in multiple studies (Reed et al. 1997; Rademakers et al. 2003; Cruts et al. 2012) (ACMG criterion PP1S).
- Functional studies (Hasegawa et al. 1998; Hong et al. 1998; Krishnamurthy and Johnson 2004; Zhang et al. 2004) (ACMG criterion PS3).
- Predicted damaging by multiple computational methods (CADD (Kircher et al. 2014), PolyPhen-2 (Adzhubei et al. 2010), and SIFT (Ng and Henikoff 2003)) (ACMG criterion PP3).
- Altogether, the presence of two strong criteria and one supporting criterion result in the ACMG-recommended assertion of “pathogenic.”

#### *CSF1R* (NM_005211.3, c.2699G>A, p.R900K)

- Critical domain of *CSF1R* where other pathogenic variants also cluster (Rademakers et al. 2011; Stabile et al. 2016) (ACMG criterion PM1)
- Absent from the gnomAD (Lek et al. 2016) and TOPMed Bravo population databases (ACMG criterion PM2)
- This particular variant has been reported before along with segregation data (Kortvelyessy et al. 2015) (ACMG criterion PP1).
- Predicted damaging by multiple computational predictions (CADD, PolyPhen-2, and SIFT) (ACMG criterion PP3).
- Taken together, the presence of two moderate criteria and two supporting criteria result in the ACMG-recommended assertion of “likely pathogenic.”

#### *PLD3* variant (NM_012268.3, c.694G>A, p.V232M)

While the *PLD3* variant described here has been controversial because of replication in some but not all cohorts tested, we considered it a “likely risk variant” based on available evidence (Cruchaga et al. 2014; Cacace et al. 2015; Cruchaga and Goate 2015b; Cruchaga and Goate 2015a; Heilmann et al. 2015; Hooli et al. 2015; Lambert et al. 2015; van der Lee et al. 2015; Engelman et al. 2018). Rare variants are not expected to replicate in all cohorts because of population effects and stochastic sampling.

#### VUS in *APP* (NM_000484.3, c.742G>A, p.D248N)

This variant (*APP* (NM_000484.3, c.742G>A, p.D248N)) was returned to the patient as a VUS, but with language indicating that, especially in the presence of the additional variants observed (*APOE* ε4 homozygosity and the *PLD3* V232M variant), it may not contribute much, if at all, to symptoms.

#### SORL1 M105T

Because this variant lies in a critical functional domain for *SORL1*, the VPS10 domain (Pottier et al. 2012; Caglayan et al. 2014; Louwersheimer et al. 2017), we computational modeled the effect of the variant. Modelling suggests this is a highly conserved residue (Fig. 3A) where change to a Threonine may create a PLK1 kinase site (Fig. 3B). PLK1 has known roles in the cell cycle, and is aberrantly present in neurons of AD patients but not age-matched controls (Song et al. 2011), leading us to speculate that presence of this variant in SORL1 may lead to faster progression of disease if this kinase phosphorylates this residue, which could disrupt the amyloid-β clearance mechanism of the VPS10 domain (Kitago et al. 2015).

- Studies where *SORL1* M105T would have been observed, but no other carriers of *SORL1* M105T were identified in either cases or controls (Vardarajan et al. 2015; Fernandez et al. 2016; Verheijen et al. 2016; Bellenguez et al. 2017).

#### TREM2

TREM2 is a well-established risk factor for AD and FTD. References for the specific variants described are as follows:

- *TREM2* (NM_018965.3, c.140G>A, p.R47H) (Guerreiro et al. 2013; Jonsson et al. 2013)
- *TREM2* (NM_018965.3, c.259G>A, p.D87N) (Guerreiro et al. 2013; Cuyvers et al. 2014; Ghani et al. 2015; Jin et al. 2015; Ghani et al. 2016; Piccio et al. 2016)

#### VPS13C Loss-of-Function Support

- unpublished studies have reported an association between heterozygous loss-of-function variant in *VPS13C* and FTD (Philtjens 2014; Picillo 2018)

### Variants of Uncertain Significance and Variants of Research Interest

Variants denoted as “Variants of Uncertain Significance” described in the following section were returned to patients because it would be possible, with limited additional information, for them to become established as associated with the patients phenotype. Variants denoted as of “research interest” in contrast were not returned to patients because it would take a great deal of evidence to establish a definitive link to the patient’s phenotype, but there is limited literature evidence indicating that it is important that we point them out to the field.

#### A possible CADASIL case with two non-Cysteine variants in NOTCH3 (D45H and G52R) spanning C49

A patient with a differential diagnosis of cerebral amyloid angiopathy, leukodystrophy, or CADASIL (Cerebral Autosomal Dominant Arteriopathy with Subcortical Infarcts and Leukoencephalopathy) (Joutel et al. 1996) harbored two variants on the same allele in *NOTCH3 (*NM_000435.2, c.133G>C, p.D45H and NM_000435.2, c.154G>A, p.G52R). While these variants do not induce a typically pathogenic alteration of a Cysteine, they do flank pathogenic variants at residue Cys49 that have been reported with three different amino acid changes (Clinvar RCV000518559.1, RCV000710993.1, RCV000518038.1, and (Joutel et al. 1996; Oki et al. 2007; Wang et al. 2011; Meng et al. 2012)). Both of the variants we observe are in ClinVar as variants of uncertain significance (RCV000518589.1 and RCV000516491.1). Furthermore, both variants are predicted damaging by CADD (27.6 and 29.5) and SIFT, and one (D45H) is predicted damaging by PolyPhen-2. We speculate that, given that these variants fall on the same haplotype, the presence of one or both of these variants may affect the function of residue Cys49 or other nearby disease-associated Cys residues such as Cys43 (Clinvar RCV000517549.1) or Cys55 (Clinvar RCV000710994.1 and RCV000516615.1). Biochemical testing for CADASIL would be informative in this case, and this haplotype was returned as a variant of uncertain significance with clear language in the report that biochemical testing should be pursued.

#### A case with a MAPT VUS

A patient with unspecified dementia with an age-of-onset in the late 40s had a VUS returned in *MAPT* (NM_005910.5, c.1174A>G, p.I392V). Family history information was incomplete for this patient, precluding knowledge of if a dominant family history was present. The variant had a CADD score of 24.6, was absent from gnomAD (out of 135,743 non-TOPMed individuals), and was present only one time in TOPMed Bravo (out of 62,784 individuals). The closest pathogenic variants are R406W (already described) and G389R (Murrell et al. 1999; Ghetti et al. 2000; Pickering-Brown et al. 2000; Bermingham et al. 2008; Rossi et al. 2008). Of note, these established pathogenic variants are present four and three times in gnomAD, respectively, indicating that the rarity of the VUS observed here justifies return to the patient as a VUS. The uncertainty around this variant was emphasized in the letter to the patient.

#### A case with APOE ε4 Heterozygosity, ADAM10 I120T, and TTC3 V1893M

A patient with corticobasal syndrome with onset in the early 50s and positive amyloid PET was found to harbor two variants of research interest, but that did not reach the level of evidence needed for return of the variants as a VUS. The variants were in *ADAM10* (NM_001110.3, c.359T>C, p.I120T) and *TTC3* (NM_001001894.2, c.5557G>A, p.V1853M). The *ADAM10* variant had a borderline CADD score of 14.3 and was not predicted damaging by PolyPhen-2 or SIFT. Furthermore, the variant was observed in gnomAD 12 times. *ADAM10* has been proposed as a candidate gene for AD in prior studies (Kim et al. 2009) including two variants in the same domain as the variant identified here, the prodomain (Suh et al. 2013). Furthermore, variation in *ADAM10* recently reached genome-wide significance for association with AD by GWAS (Marioni et al. 2018; Kunkle et al. 2019). However, we have chosen to not return this variant in the absence of more information about effect size or segregation. The *TTC3* variant also had a borderline CADD score (14.6) and was not predicted damaging by PolyPhen-2 or SIFT. However, this variant was not observed in gnomAD or TOPMed Bravo. A different *TTC3* variant (NM_001001894.2, c.3113C>G, p.S1038C) has been reported to segregate with late-onset AD in one family (Kohli et al. 2016). However, since we lacked segregation data for the variant we observed, we did not have enough evidence to consider the *TTC3* variant as more than a variant of research interest, and thus did not return the variant to the patient.

#### A case with APOE ε4 Heterozygosity, SORL1 R416Q, and MME Y414C

A case with mild dementia of uncertain etiology and symptoms consistent with neuropathy with onset in the mid 50s had one *APOE* ε4 allele along with variants in *SORL1* (NM_003105.5, c.1247G>A, p.R416Q) and *MME* (NM_007289.2, c.1241A>G, p.Y414C). This *SORL1* variant has a CADD score of 34 and is also predicted damaging by PolyPhen-2 and SIFT. A link between *MME* and neurodegeneration, including AD and neuropathy, has previously been proposed (Rey-Salgueiro et al. 2009; Auer-Grumbach et al. 2016; Depondt et al. 2016), but there was insufficient evidence for this particular variant in *MME* or for the *SORL1* variant to justify return to the patient.

#### A case with TM2D3 P69L and TNK1 H131Q

A patient with mild dementia due to either AD or bvFTD with onset in the mid 50s had variants in *TM2D3* (NM_078474.2, c.206C>T, p.P69L) and *TNK1* (NM_001251902.1, c.393C>G, p.H131Q). A different variant in *TM2D3* has been nominated as AD-associated from an Icelandic cohort (Jakobsdottir et al. 2016). Other variants in *TNK1* have been nominated as AD-associated from analysis of Alzheimer’s Disease Sequencing Project data (He et al. 2017). While neither of these variants had sufficient evidence for return as risk variants, our observation of these variants in this cohort adds evidence for the possible contribution of variants in these genes to disease.

#### Secondary Finding

One patient harbored a secondary pathogenic variant in *KCNQ1* (NM_000218.2, c.1552C>T, R518*), associated with cardiac arrhythmias. This is a known founder effect variant from the Swedish population that responds well to beta blockers (Winbo et al. 2014). The variant is a null variant in a gene where loss-of-function is a known mechanism of disease (ACMG criterion PVS1) and is enriched in cases vs. controls with an odds ratio >5 (ACMG criterion PS4) (Kapplinger et al. 2009). Furthermore, the variant’s effect is supported by well-established functional studies (Harmer et al. 2014) (ACMG criterion PS3). Taken together, the presence of one very strong criterion and two strong criteria result in the ACMG-recommended assertion of “pathogenic.” Consistent with the study consent and protocol, presence of this variant was reported to the patient.

## Extended Acknowledgements

### ADSP

The Alzheimer’s Disease Sequencing Project (ADSP) is comprised of two Alzheimer’s Disease (AD) genetics consortia and three National Human Genome Research Institute (NHGRI) funded Large Scale Sequencing and Analysis Centers (LSAC). The two AD genetics consortia are the Alzheimer’s Disease Genetics Consortium (ADGC) funded by NIA (U01 AG032984), and the Cohorts for Heart and Aging Research in Genomic Epidemiology (CHARGE) funded by NIA (R01 AG033193), the National Heart, Lung, and Blood Institute (NHLBI), other National Institute of Health (NIH) institutes and other foreign governmental and non-governmental organizations. The Discovery Phase analysis of sequence data is supported through UF1AG047133 (to Drs. Schellenberg, Farrer, Pericak-Vance, Mayeux, and Haines); U01AG049505 to Dr. Seshadri; U01AG049506 to Dr. Boerwinkle; U01AG049507 to Dr. Wijsman; and U01AG049508 to Dr. Goate and the Discovery Extension Phase analysis is supported through U01AG052411 to Dr. Goate, U01AG052410 to Dr. Pericak-Vance and U01 AG052409 to Drs. Seshadri and Fornage. Data generation and harmonization in the Follow-up Phases is supported by U54AG052427 (to Drs. Schellenberg and Wang).

The ADGC cohorts include: Adult Changes in Thought (ACT), the Alzheimer’s Disease Centers (ADC), the Chicago Health and Aging Project (CHAP), the Memory and Aging Project (MAP), Mayo Clinic (MAYO), Mayo Parkinson’s Disease controls, University of Miami, the Multi-Institutional Research in Alzheimer’s Genetic Epidemiology Study (MIRAGE), the National Cell Repository for Alzheimer’s Disease (NCRAD), the National Institute on Aging Late Onset Alzheimer’s Disease Family Study (NIA-LOAD), the Religious Orders Study (ROS), the Texas Alzheimer’s Research and Care Consortium (TARC), Vanderbilt University/Case Western Reserve University (VAN/CWRU), the Washington Heights-Inwood Columbia Aging Project (WHICAP) and the Washington University Sequencing Project (WUSP), the Columbia University Hispanic-Estudio Familiar de Influencia Genetica de Alzheimer (EFIGA), the University of Toronto (UT), and Genetic Differences (GD).

The CHARGE cohorts are supported in part by National Heart, Lung, and Blood Institute (NHLBI) infrastructure grant HL105756 (Psaty), RC2HL102419 (Boerwinkle) and the neurology working group is supported by the National Institute on Aging (NIA) R01 grant AG033193. The CHARGE cohorts participating in the ADSP include the following: Austrian Stroke Prevention Study (ASPS), ASPS-Family study, and the Prospective Dementia Registry-Austria (ASPS/PRODEM-Aus), the Atherosclerosis Risk in Communities (ARIC) Study, the Cardiovascular Health Study (CHS), the Erasmus Rucphen Family Study (ERF), the Framingham Heart Study (FHS), and the Rotterdam Study (RS). ASPS is funded by the Austrian Science Fond (FWF) grant number P20545-P05 and P13180 and the Medical University of Graz. The ASPS-Fam is funded by the Austrian Science Fund (FWF) project I904),the EU Joint Programme-Neurodegenerative Disease Research (JPND) in frame of the BRIDGET project (Austria, Ministry of Science) and the Medical University of Graz and the Steiermärkische Krankenanstalten Gesellschaft. PRODEM-Austria is supported by the Austrian Research Promotion agency (FFG) (Project No. 827462) and by the Austrian National Bank (Anniversary Fund, project 15435. ARIC research is carried out as a collaborative study supported by NHLBI contracts (HHSN268201100005C, HHSN268201100006C, HHSN268201100007C, HHSN268201100008C, HHSN268201100009C, HHSN268201100010C, HHSN268201100011C, and HHSN268201100012C). Neurocognitive data in ARIC is collected by U01 2U01HL096812, 2U01HL096814, 2U01HL096899, 2U01HL096902, 2U01HL096917 from the NIH (NHLBI, NINDS, NIA and NIDCD), and with previous brain MRI examinations funded by R01-HL70825 from the NHLBI. CHS research was supported by contracts HHSN268201200036C, HHSN268200800007C, N01HC55222, N01HC85079, N01HC85080, N01HC85081, N01HC85082, N01HC85083, N01HC85086, and grants U01HL080295 and U01HL130114 from the NHLBI with additional contribution from the National Institute of Neurological Disorders and Stroke (NINDS). Additional support was provided by R01AG023629, R01AG15928, and R01AG20098 from the NIA. FHS research is supported by NHLBI contracts N01-HC-25195 and HHSN268201500001I. This study was also supported by additional grants from the NIA (R01s AG054076, AG049607 and AG033040 and NINDS (R01 NS017950). The ERF study as a part of EUROSPAN (European Special Populations Research Network) was supported by European Commission FP6 STRP grant number 018947 (LSHG-CT-2006-01947) and also received funding from the European Community’s Seventh Framework Programme (FP7/2007-2013)/grant agreement HEALTH-F4-2007-201413 by the European Commission under the programme “Quality of Life and Management of the Living Resources” of 5th Framework Programme (no. QLG2-CT-2002-01254). High-throughput analysis of the ERF data was supported by a joint grant from the Netherlands Organization for Scientific Research and the Russian Foundation for Basic Research (NWO-RFBR 047.017.043). The Rotterdam Study is funded by Erasmus Medical Center and Erasmus University, Rotterdam, the Netherlands Organization for Health Research and Development (ZonMw), the Research Institute for Diseases in the Elderly (RIDE), the Ministry of Education, Culture and Science, the Ministry for Health, Welfare and Sports, the European Commission (DG XII), and the municipality of Rotterdam. Genetic data sets are also supported by the Netherlands Organization of Scientific Research NWO Investments (175.010.2005.011, 911-03-012), the Genetic Laboratory of the Department of Internal Medicine, Erasmus MC, the Research Institute for Diseases in the Elderly (014-93-015; RIDE2), and the Netherlands Genomics Initiative (NGI)/Netherlands Organization for Scientific Research (NWO) Netherlands Consortium for Healthy Aging (NCHA), project 050-060-810. All studies are grateful to their participants, faculty and staff. The content of these manuscripts is solely the responsibility of the authors and does not necessarily represent the official views of the National Institutes of Health or the U.S. Department of Health and Human Services.

The four LSACs are: the Human Genome Sequencing Center at the Baylor College of Medicine (U54 HG003273), the Broad Institute Genome Center (U54HG003067), The American Genome Center at the Uniformed Services University of the Health Sciences (U01AG057659), and the Washington University Genome Institute (U54HG003079).

Biological samples and associated phenotypic data used in primary data analyses were stored at Study Investigators institutions, and at the National Cell Repository for Alzheimer’s Disease (NCRAD, U24AG021886) at Indiana University funded by NIA. Associated Phenotypic Data used in primary and secondary data analyses were provided by Study Investigators, the NIA funded Alzheimer’s Disease Centers (ADCs), and the National Alzheimer’s Coordinating Center (NACC, U01AG016976) and the National Institute on Aging Genetics of Alzheimer’s Disease Data Storage Site (NIAGADS, U24AG041689) at the University of Pennsylvania, funded by NIA, and at the Database for Genotypes and Phenotypes (dbGaP) funded by NIH. This research was supported in part by the Intramural Research Program of the National Institutes of health, National Library of Medicine. Contributors to the Genetic Analysis Data included Study Investigators on projects that were individually funded by NIA, and other NIH institutes, and by private U.S. organizations, or foreign governmental or nongovernmental organizations.

### ADNI

Michael Weiner, MD (UC San Francisco, Principal Investigator, Executive Committee); Paul Aisen, MD (UC San Diego, ADCS PI and Director of Coordinating Center Clinical Core, Executive Committee, Clinical Core Leaders); Ronald Petersen, MD, PhD (Mayo Clinic, Rochester, Executive Committee, Clinical Core Leader); Clifford R. Jack, Jr., MD (Mayo Clinic, Rochester, Executive Committee, MRI Core Leader); William Jagust, MD (UC Berkeley, Executive Committee; PET Core Leader); John Q. Trojanowki, MD, PhD (U Pennsylvania, Executive Committee, Biomarkers Core Leader); Arthur W. Toga, PhD (USC, Executive Committee, Informatics Core Leader); Laurel Beckett, PhD (UC Davis, Executive Committee, Biostatistics Core Leader); Robert C. Green, MD, MPH (Brigham and Women’s Hospital, Harvard Medical School, Executive Committee and Chair of Data and Publication Committee); Andrew J. Saykin, PsyD (Indiana University, Executive Committee, Genetics Core Leader); John Morris, MD (Washington University St. Louis, Executive Committee, Neuropathology Core Leader); Leslie M. Shaw (University of Pennsylvania, Executive Committee, Biomarkers Core Leader); Enchi Liu, PhD (Janssen Alzheimer Immunotherapy, ADNI two Private Partner Scientific Board Chair); Tom Montine, MD, PhD (University of Washington); Ronald G. Thomas, PhD (UC San Diego); Michael Donohue, PhD (UC San Diego); Sarah Walter, MSc (UC San Diego); Devon Gessert (UC San Diego); Tamie Sather, MS (UC San Diego,); Gus Jiminez, MBS (UC San Diego); Danielle Harvey, PhD (UC Davis;); Michael Donohue, PhD (UC San Diego); Matthew Bernstein, PhD (Mayo Clinic, Rochester); Nick Fox, MD (University of London); Paul Thompson, PhD (USC School of Medicine); Norbert Schuff, PhD (UCSF MRI); Charles DeCArli, MD (UC Davis); Bret Borowski, RT (Mayo Clinic); Jeff Gunter, PhD (Mayo Clinic); Matt Senjem, MS (Mayo Clinic); Prashanthi Vemuri, PhD (Mayo Clinic); David Jones, MD (Mayo Clinic); Kejal Kantarci (Mayo Clinic); Chad Ward (Mayo Clinic); Robert A. Koeppe, PhD (University of Michigan, PET Core Leader); Norm Foster, MD (University of Utah); Eric M. Reiman, MD (Banner Alzheimer’s Institute); Kewei Chen, PhD (Banner Alzheimer’s Institute); Chet Mathis, MD (University of Pittsburgh); Susan Landau, PhD (UC Berkeley); Nigel J. Cairns, PhD, MRCPath (Washington University St. Louis); Erin Householder (Washington University St. Louis); Lisa Taylor Reinwald, BA, HTL (Washington University St. Louis); Virginia Lee, PhD, MBA (UPenn School of Medicine); Magdalena Korecka, PhD (UPenn School of Medicine); Michal Figurski, PhD (UPenn School of Medicine); Karen Crawford (USC); Scott Neu, PhD (USC); Tatiana M. Foroud, PhD (Indiana University); Steven Potkin, MD UC (UC Irvine); Li Shen, PhD (Indiana University); Faber Kelley, MS, CCRC (Indiana University); Sungeun Kim, PhD (Indiana University); Kwangsik Nho, PhD (Indiana University); Zaven Kachaturian, PhD (Khachaturian, Radebaugh & Associates, Inc and Alzheimer’s Association’s Ronald and Nancy Reagan’s Research Institute); Richard Frank, MD, PhD (General Electric); Peter J. Snyder, PhD (Brown University); Susan Molchan, PhD (National Institute on Aging/ National Institutes of Health); Jeffrey Kaye, MD (Oregon Health and Science University); Joseph Quinn, MD (Oregon Health and Science University); Betty Lind, BS (Oregon Health and Science University); Raina Carter, BA (Oregon Health and Science University); Sara Dolen, BS (Oregon Health and Science University); Lon S. Schneider, MD (University of Southern California); Sonia Pawluczyk, MD (University of Southern California); Mauricio Beccera, BS (University of Southern California); Liberty Teodoro, RN (University of Southern California); Bryan M. Spann, DO, PhD (University of Southern California); James Brewer, MD, PhD (University of California San Diego); Helen Vanderswag, RN (University of California San Diego); Adam Fleisher, MD (University of California San Diego); Judith L. Heidebrink, MD, MS (University of Michigan); Joanne L. Lord, LPN, BA, CCRC (University of Michigan); Ronald Petersen, MD, PhD (Mayo Clinic, Rochester); Sara S. Mason, RN (Mayo Clinic, Rochester); Colleen S. Albers, RN (Mayo Clinic, Rochester); David Knopman, MD (Mayo Clinic, Rochester); Kris Johnson, RN (Mayo Clinic, Rochester); Rachelle S. Doody, MD, PhD (Baylor College of Medicine); Javier Villanueva Meyer, MD (Baylor College of Medicine); Munir Chowdhury, MBBS, MS (Baylor College of Medicine); Susan Rountree, MD (Baylor College of Medicine); Mimi Dang, MD (Baylor College of Medicine); Yaakov Stern, PhD (Columbia University Medical Center); Lawrence S. Honig, MD, PhD (Columbia University Medical Center); Karen L. Bell, MD (Columbia University Medical Center); Beau Ances, MD (Washington University, St. Louis); John C. Morris, MD (Washington University, St. Louis); Maria Carroll, RN, MSN (Washington University, St. Louis); Sue Leon, RN, MSN (Washington University, St. Louis); Erin Householder, MS, CCRP (Washington University, St. Louis); Mark A. Mintun, MD (Washington University, St. Louis); Stacy Schneider, APRN, BC, GNP (Washington University, St. Louis); Angela Oliver, RN, BSN, MSG; Daniel Marson, JD, PhD (University of Alabama Birmingham); Randall Griffith, PhD, ABPP (University of Alabama Birmingham); David Clark, MD (University of Alabama Birmingham); David Geldmacher, MD (University of Alabama Birmingham); John Brockington, MD (University of Alabama Birmingham); Erik Roberson, MD (University of Alabama Birmingham); Hillel Grossman, MD (Mount Sinai School of Medicine); Effie Mitsis, PhD (Mount Sinai School of Medicine); Leyla deToledo-Morrell, PhD (Rush University Medical Center); Raj C. Shah, MD (Rush University Medical Center); Ranjan Duara, MD (Wien Center); Daniel Varon, MD (Wien Center); Maria T. Greig, HP (Wien Center); Peggy Roberts, CNA (Wien Center); Marilyn Albert, PhD (Johns Hopkins University); Chiadi Onyike, MD (Johns Hopkins University); Daniel D’Agostino II, BS (Johns Hopkins University); Stephanie Kielb, BS (Johns Hopkins University); James E. Galvin, MD, MPH (New York University); Dana M. Pogorelec (New York University); Brittany Cerbone (New York University); Christina A. Michel (New York University); Henry Rusinek, PhD (New York University); Mony J de Leon, EdD (New York University); Lidia Glodzik, MD, PhD (New York University); Susan De Santi, PhD (New York University); P. Murali Doraiswamy, MD (Duke University Medical Center); Jeffrey R. Petrella, MD (Duke University Medical Center); Terence Z. Wong, MD (Duke University Medical Center); Steven E. Arnold, MD (University of Pennsylvania); Jason H. Karlawish, MD (University of Pennsylvania); David Wolk, MD (University of Pennsylvania); Charles D. Smith, MD (University of Kentucky); Greg Jicha, MD (University of Kentucky); Peter Hardy, PhD (University of Kentucky); Partha Sinha, PhD (University of Kentucky); Elizabeth Oates, MD (University of Kentucky); Gary Conrad, MD (University of Kentucky); Oscar L. Lopez, MD (University of Pittsburgh); MaryAnn Oakley, MA (University of Pittsburgh); Donna M. Simpson, CRNP, MPH (University of Pittsburgh); Anton P. Porsteinsson, MD (University of Rochester Medical Center); Bonnie S. Goldstein, MS, NP (University of Rochester Medical Center); Kim Martin, RN (University of Rochester Medical Center); Kelly M. Makino, BS (University of Rochester Medical Center); M. Saleem Ismail, MD (University of Rochester Medical Center); Connie Brand, RN (University of Rochester Medical Center); Ruth A. Mulnard, DNSc, RN, FAAN (University of California, Irvine); Gaby Thai, MD (University of California, Irvine); Catherine Mc Adams Ortiz, MSN, RN, A/GNP (University of California, Irvine); Kyle Womack, MD (University of Texas Southwestern Medical School); Dana Mathews, MD, PhD (University of Texas Southwestern Medical School); Mary Quiceno, MD (University of Texas Southwestern Medical School); Ramon Diaz Arrastia, MD, PhD (University of Texas Southwestern Medical School); Richard King, MD (University of Texas Southwestern Medical School); Myron Weiner, MD (University of Texas Southwestern Medical School); Kristen Martin Cook, MA (University of Texas Southwestern Medical School); Michael DeVous, PhD (University of Texas Southwestern Medical School); Allan I. Levey, MD, PhD (Emory University); James J. Lah, MD, PhD (Emory University); Janet S. Cellar, DNP, PMHCNS BC (Emory University); Jeffrey M. Burns, MD (University of Kansas, Medical Center); Heather S. Anderson, MD (University of Kansas, Medical Center); Russell H. Swerdlow, MD (University of Kansas, Medical Center); Liana Apostolova, MD (University of California, Los Angeles); Kathleen Tingus, PhD (University of California, Los Angeles); Ellen Woo, PhD (University of California, Los Angeles); Daniel H.S. Silverman, MD, PhD (University of California, Los Angeles); Po H. Lu, PsyD (University of California, Los Angeles); George Bartzokis, MD (University of California, Los Angeles); Neill R Graff Radford, MBBCH, FRCP (London) (Mayo Clinic, Jacksonville); Francine Parfitt, MSH, CCRC (Mayo Clinic, Jacksonville); Tracy Kendall, BA, CCRP (Mayo Clinic, Jacksonville); Heather Johnson, MLS, CCRP (Mayo Clinic, Jacksonville); Martin R. Farlow, MD (Indiana University); Ann Marie Hake, MD (Indiana University); Brandy R. Matthews, MD (Indiana University); Scott Herring, RN, CCRC (Indiana University); Cynthia Hunt, BS, CCRP (Indiana University); Christopher H. van Dyck, MD (Yale University School of Medicine); Richard E. Carson, PhD (Yale University School of Medicine); Martha G. MacAvoy, PhD (Yale University School of Medicine); Howard Chertkow, MD (McGill Univ., Montreal Jewish General Hospital); Howard Bergman, MD (McGill Univ., Montreal Jewish General Hospital); Chris Hosein, Med (McGill Univ., Montreal Jewish General Hospital); Sandra Black, MD, FRCPC (Sunnybrook Health Sciences, Ontario); Dr Bojana Stefanovic (Sunnybrook Health Sciences, Ontario); Curtis Caldwell, PhD (Sunnybrook Health Sciences, Ontario); Ging Yuek Robin Hsiung, MD, MHSc, FRCPC (U.B.C. Clinic for AD & Related Disorders); Howard Feldman, MD, FRCPC (U.B.C. Clinic for AD & Related Disorders); Benita Mudge, BS (U.B.C. Clinic for AD & Related Disorders); Michele Assaly, MA Past (U.B.C. Clinic for AD & Related Disorders); Andrew Kertesz, MD (Cognitive Neurology St. Joseph’s, Ontario); John Rogers, MD (Cognitive Neurology St. Joseph’s, Ontario); Dick Trost, PhD (Cognitive Neurology St. Joseph’s, Ontario); Charles Bernick, MD (Cleveland Clinic Lou Ruvo Center for Brain Health); Donna Munic, PhD (Cleveland Clinic Lou Ruvo Center for Brain Health); Diana Kerwin, MD (Northwestern University); Marek Marsel Mesulam, MD (Northwestern University); Kristine Lipowski, BA (Northwestern University); Chuang Kuo Wu, MD, PhD (Northwestern University); Nancy Johnson, PhD (Northwestern University); Carl Sadowsky, MD (Premiere Research Inst (Palm Beach Neurology)); Walter Martinez, MD (Premiere Research Inst (Palm Beach Neurology)); Teresa Villena, MD (Premiere Research Inst (Palm Beach Neurology)); Raymond Scott Turner, MD, PhD (Georgetown University Medical Center); Kathleen Johnson, NP (Georgetown University Medical Center); Brigid Reynolds, NP (Georgetown University Medical Center); Reisa A. Sperling, MD (Brigham and Women’s Hospital); Keith A. Johnson, MD (Brigham and Women’s Hospital); Gad Marshall, MD (Brigham and Women’s Hospital); Meghan Frey (Brigham and Women’s Hospital); Jerome Yesavage, MD (Stanford University); Joy L. Taylor, PhD (Stanford University); Barton Lane, MD (Stanford University); Allyson Rosen, PhD (Stanford University); Jared Tinklenberg, MD (Stanford University); Marwan N. Sabbagh, MD (Banner Sun Health Research Institute); Christine M. Belden, PsyD (Banner Sun Health Research Institute); Sandra A. Jacobson, MD (Banner Sun Health Research Institute); Sherye A. Sirrel, MS (Banner Sun Health Research Institute); Neil Kowall, MD (Boston University); Ronald Killiany, PhD (Boston University); Andrew E. Budson, MD (Boston University); Alexander Norbash, MD (Boston University); Patricia Lynn Johnson, BA (Boston University); Thomas O. Obisesan, MD, MPH (Howard University); Saba Wolday, MSc (Howard University); Joanne Allard, PhD (Howard University); Alan Lerner, MD (Case Western Reserve University); Paula Ogrocki, PhD (Case Western Reserve University); Leon Hudson, MPH (Case Western Reserve University); Evan Fletcher, PhD (University of California, Davis Sacramento); Owen Carmichael, PhD (University of California, Davis Sacramento); John Olichney, MD (University of California, Davis Sacramento); Charles DeCarli, MD (University of California, Davis Sacramento); Smita Kittur, MD (Neurological Care of CNY); Michael Borrie, MB ChB (Parkwood Hospital); T Y Lee, PhD (Parkwood Hospital); Dr Rob Bartha, PhD (Parkwood Hospital); Sterling Johnson, PhD (University of Wisconsin); Sanjay Asthana, MD (University of Wisconsin); Cynthia M. Carlsson, MD (University of Wisconsin); Steven G. Potkin, MD (University of California, Irvine BIC); Adrian Preda, MD (University of California, Irvine BIC); Dana Nguyen, PhD (University of California, Irvine BIC); Pierre Tariot, MD (Banner Alzheimer’s Institute); Adam Fleisher, MD (Banner Alzheimer’s Institute); Stephanie Reeder, BA (Banner Alzheimer’s Institute); Vernice Bates, MD (Dent Neurologic Institute); Horacio Capote, MD (Dent Neurologic Institute); Michelle Rainka, PharmD, CCRP (Dent Neurologic Institute); Douglas W. Scharre, MD (Ohio State University); Maria Kataki, MD, PhD (Ohio State University); Anahita Adeli, MD (Ohio State University); Earl A. Zimmerman, MD (Albany Medical College); Dzintra Celmins, MD (Albany Medical College); Alice D. Brown, FNP (Albany Medical College); Godfrey D. Pearlson, MD (Hartford Hosp, Olin Neuropsychiatry Research Center); Karen Blank, MD (Hartford Hosp, Olin Neuropsychiatry Research Center); Karen Anderson, RN (Hartford Hosp, Olin Neuropsychiatry Research Center); Robert B. Santulli, MD (Dartmouth Hitchcock Medical Center); Tamar J. Kitzmiller (Dartmouth Hitchcock Medical Center); Eben S. Schwartz, PhD (Dartmouth Hitchcock Medical Center); Kaycee M. Sink, MD, MAS (Wake Forest University Health Sciences); Jeff D. Williamson, MD, MHS (Wake Forest University Health Sciences); Pradeep Garg, PhD (Wake Forest University Health Sciences); Franklin Watkins, MD (Wake Forest University Health Sciences); Brian R. Ott, MD (Rhode Island Hospital); Henry Querfurth, MD (Rhode Island Hospital); Geoffrey Tremont, PhD (Rhode Island Hospital); Stephen Salloway, MD, MS (Butler Hospital); Paul Malloy, PhD (Butler Hospital); Stephen Correia, PhD (Butler Hospital); Howard J. Rosen, MD (UC San Francisco); Bruce L. Miller, MD (UC San Francisco); Jacobo Mintzer, MD, MBA (Medical University South Carolina); Kenneth Spicer, MD, PhD (Medical University South Carolina); David Bachman, MD (Medical University South Carolina); Elizabether Finger, MD (St. Joseph’s Health Care); Stephen Pasternak, MD (St. Joseph’s Health Care); Irina Rachinsky, MD (St. Joseph’s Health Care); John Rogers, MD (St. Joseph’s Health Care); Andrew Kertesz, MD (St. Joseph’s Health Care); Dick Drost, MD (St. Joseph’s Health Care); Nunzio Pomara, MD (Nathan Kline Institute); Raymundo Hernando, MD (Nathan Kline Institute); Antero Sarrael, MD (Nathan Kline Institute); Susan K. Schultz, MD (University of Iowa College of Medicine, Iowa City); Laura L. Boles Ponto, PhD (University of Iowa College of Medicine, Iowa City); Hyungsub Shim, MD (University of Iowa College of Medicine, Iowa City); Karen Elizabeth Smith, RN (University of Iowa College of Medicine, Iowa City); Norman Relkin, MD, PhD (Cornell University); Gloria Chaing, MD (Cornell University); Lisa Raudin, PhD (Cornell University); Amanda Smith, MD (University of South Floriday: USF Health Byrd Alzheimer’s Institute); Kristin Fargher, MD (University of South Floriday: USF Health Byrd Alzheimer’s Institute); Balebail Ashok Raj, MD (University of South Floriday: USF Health Byrd Alzheimer’s Institute)

## FIGURE LEGENDS

**Supplemental Table 1:** Phenotype and variant table. Prior clinical diagnosis category, age of onset range, family history score, **Figure 1** category, and variant information listed in **Table 1** for each proband are listed along with information on which variants were returned to patients and which did not have sufficient evidence for return but are of research interest. Note that some detailed information such as sex, age of onset to the year, self-reported ethnicity, and detailed phenotype and family history information has been excluded to protect the identity of participants but is available along with raw data via controlled access to qualified researchers.

